# Gene age shapes functional and evolutionary properties of the *Drosophila* seminal fluid proteome

**DOI:** 10.1101/2024.12.17.628492

**Authors:** Jose M. Ranz, Carolina Flacchi, Imtiyaz E. Hariyani, Alberto Civetta

**Author notes:** **Correspondence and requests for materials should be addressed to:** Jose Ranz and Alberto Civetta.

## Abstract

Seminal fluid proteins (Sfps) are crucial for animal reproductive success, with most Sfps-encoding genes believed to be evolutionarily young and rapidly evolving. By employing estimates of the phylogenetic origin of each *Drosophila melanogaster* Sfp gene based on genomic resources that include outgroup species to the *Drosophila* genus, we examined the functional attributes and evolutionary characteristics of Sfp genes relative to their evolutionary age. Contrary to common belief, 62% of Sfp genes existed in the genome of the ancestor to such genus. These ancient genes have broadened their expression profiles, expanded their biological roles, and formed a denser network of interactions with non-Sfp genes. This increased pleiotropy has imposed constraints on the rate of sequence evolution in ancient Sfp genes compared to younger ones. Within the Sfp interactome, we identified a fast-evolving core sub-network of younger genes with more restricted tissue expression and functions. Our findings uncover a large, previously unrecognized set of ancient Sfp genes with distinct genomic, functional, and evolutionary characteristics compared to the younger, more commonly studied Sfp genes.

## Introduction

Seminal fluid proteins (Sfps) are key components of the male ejaculate that is transferred along with the sperm to the female during copulation. Sfps play central roles in postmating female processes, affecting individual fitness ^1-4^. In *Drosophila*, comparative genomic analyses have shown a dynamic Sfp gene complement, with many genes evolving rapidly in sequence and expression, and experiencing high rates of duplication and loss ^5-9^. However, no thorough analysis has clarified the origins of the entire *Drosophila* Sfp gene complement or assessed whether gene age affects their evolutionary properties. It also remains unknown whether the expression and function of ancient Sfp genes is primarily linked to reproduction. Additionally, the evolution of Sfps as a network of interactive partners remains uncharacterized.

Previous studies on the network topology and evolution of the *D. melanogaster* Sfp genes have been focused on the Sfp sex peptide (SP) due to its role in female postmating responses ^10-12^. SP binds to the sperm, which is facilitated by a network of at least eight additional Sfps that assist in sperm storage ^10,12^. The SP network genes were inferred to have originated before the evolution of SP-mediated responses in females, with both purifying and positive selection shaping their sequence evolution ^11^. However, an analysis of the entire Sfp network, considering the incorporation of new Sfp genes into the genome over evolutionary time, is lacking. This gap prevents a full understanding of the evolution of the seminal fluid proteome function, one that recognizes polygenic phenotypic outputs, transitioning from single gene studies to systems biology ^13^.

Here, we delineated a robust catalog of *D. melanogaster* Sfp-encoding genes based on high-confidence calls from two independent research groups ^14-16^, and characterized the functional and evolutionary properties of Sfp genes. We do so while exploring how these properties relate to the time of origin of these genes within the *Drosophila* lineage and more distant phylogenetic timepoints ^17^. Additionally, we analyzed Sfps as an evolving protein network. Our findings offer significant insights into how Sfp genes have evolved in terms of their interactions and evolutionary age. Contrary to the predominant perception ^5-9^, we find that a large portion of Sfp genes in *D. melanogaster* originated prior to the diversification of the genus *Drosophila*. These ancient Sfp genes are peripheral to the network, show pleiotropic expression and functionality, frequently interact outside the Sfp network, and exhibit slower nucleotide change rates. In contrast, younger Sfp genes dominate the core network, are enriched in reproductive functions, and show faster rates of change.

## RESULTS AND DISCUSSION

### The Sfp gene complement is evolutionarily ancient

We cataloged 357 Sfp-encoding genes in *D. melanogaster*, combining results from two research groups using partially different criteria (Methods). Approximately two-thirds (228/357) of the Sfp gene candidates were common to both studies, with the rest equally contributed by each (Fig. 1a, Supplementary Table 1).

**Figure 1.**
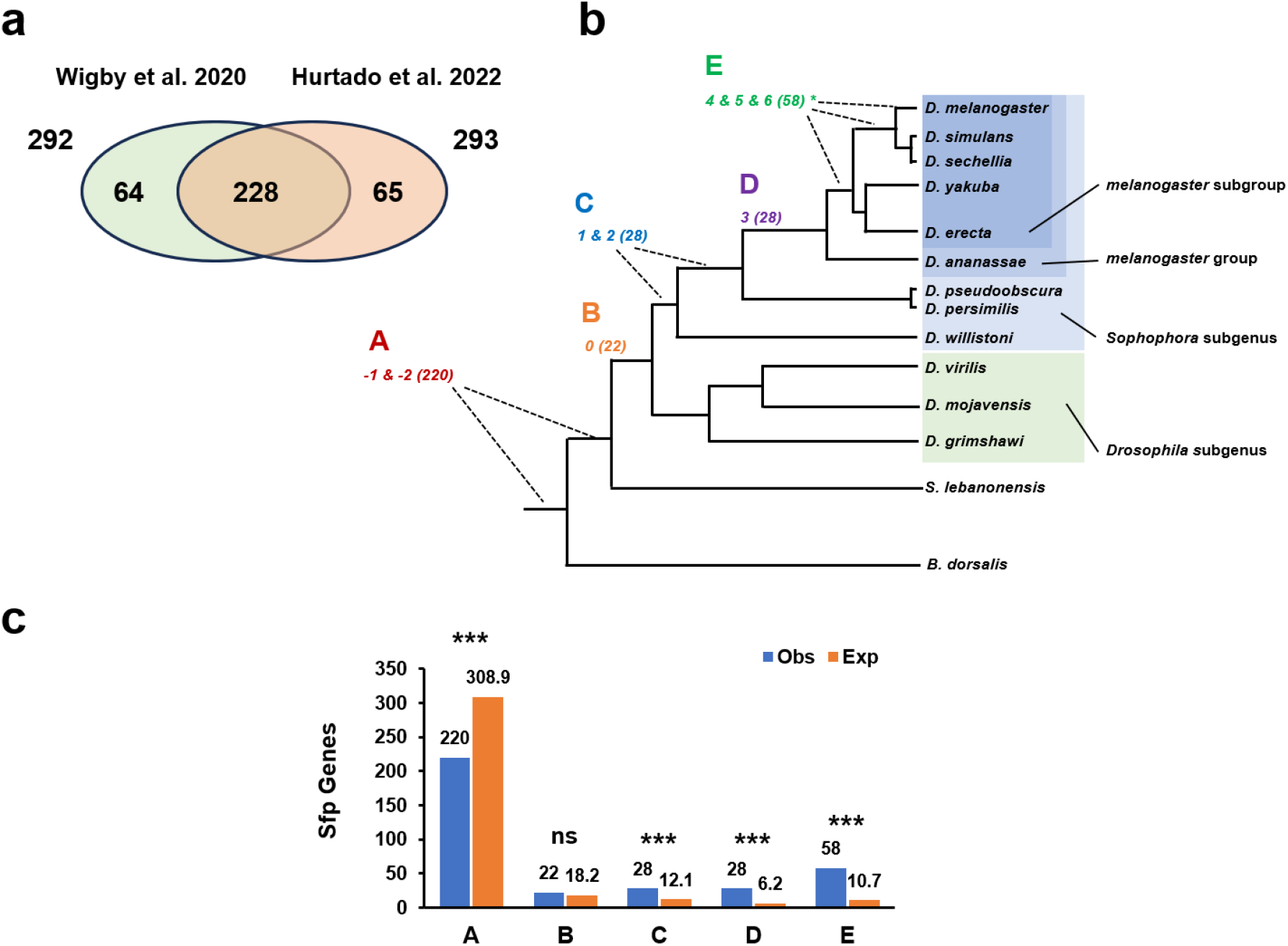
Age categorization of *D. melanogaster* Sfp-encoding genes. **a** Relationship of the datasets used to create a catalog of 357 high-confidence Sfp genes. **b** Phylogenetic tree of key species used to infer the origin of Sfp-encoding genes ^18,19^, with five gene age classes corresponding to the branch codes used by Dong et al. ^17^: class A, genes present before the *Drosophila* radiation (branches -1 and -2); class B, genes originated before the split between the *Drosophila* and *Sophophora* subgenera (branch 0); class C, genes formed in early divergent lineages leading to *D. willistoni* and *D. pseudoobscura* species groups (branches 1 and 2); class D, genes originated in the melanogaster species group (branch 3); and class E, genes present only in the *D. melanogaster* species subgroup (branches 4-6), including the simulans species complex and *D. melanogaster*. The number of Sfp genes originated within each age class is indicated in parentheses. **c** Clustered column plot showing the observed (blue) and expected (orange) counts of Sfp-encoding genes across age classes. Expected counts were calculated based on the proportion of these age classes in the entire *D. melanogaster* gene complement ^17^. The asterisks indicate particular age classes for which the difference between observed and expected counts are statistically significant according to the standardized residual values of the chi-square test of independence: *, <0.05; **, <0.01; ***, <0.001 ^24^.

We then estimated each gene’s origin within the species phylogeny using gene ages inferred via a maximum parsimonious framework based on multiple sequence alignments as well as microsynteny from multiple fly species ^17^. Two species, *Scaptodrosophila lebanonensis* and *Bactrocera dorsalis*, serve as increasingly distant outgroups to the genus *Drosophila* ^18,19^, providing essential phylogenetic depth for reliable gene age estimation.

All genes but one (LysC, a pseudogene) were categorized into one of five age classes spanning key sections of the species phylogeny (A-E from oldest to youngest; Fig. 1b, Supplementary Table 1). We validated our gene age classification using independent data. Specifically, we used information from 90 *D. pseudoobscura* proteins with 1-to-1 pairwise orthology to known *D. melanogaster* Sfps ^20^, plus 133 gene models from an upgraded annotation of *D. willistoni*, also 1-to-1 orthologs with *D. melanogaster* Sfp genes ^21^. In total, we cross-checked 178 *D. melanogaster* Sfp genes (Supplementary Table 1) that should belong to age classes A, B, or C, and found only five discrepancies (*Acp26Aa, Sfp33A1, Sfp53D, CG34051*, and *CG4271*), *i*.*e*. genes assigned to younger age classes (D and E). These discrepancies might result from errors in the gene age assignments, misidentifications in proteomic assays, or orthology call inaccuracies between *D. melanogaster* and *D. willistoni*. The fact that only 2.81% (5/178) of the checked Sfp genes showed discrepancies supports the reliability of the gene age classification adopted.

Two-hundred and twenty Sfp genes originated prior to, and 136 during, the radiation of the *Drosophila* genus (class A vs classes B-E: 62.0% vs 38.0%; Fig. 1b and Supplementary Fig. 1a). However, age class A genes are underrepresented compared to their share in the entire genome (chi-square goodness-of-fit, χ^2^=333.74, d.f.=4, *P*<2.2×10^-16^) (Fig. 1c, Supplementary Table 2). This conclusion held when using the more conservative set of 228 Sfp genes (chi-square goodness-of-fit, χ^2^=333.71, d.f.=4, *P*<2.2×10^-16^) (Supplementary Fig. 1a and Supplementary Table 2). Notably, these results emphasize the ancient origins of many Sfp gene members, contradicting the relatively recent origin commonly attributed to Sfp genes ^5,6,8^. This pattern is exemplified by the nine known genes of the SP network ^10,22,23^. Of them, eight genes are part of age class A (*antr, aqrs, CG9997, CG17575, intr, lectin-46Ca, lectin-46Cb* and *SP*), and one is part of age class B (*Sems*), meaning all were present prior the diversification of the genus *Drosophila*.

### Ancient Sfp genes are more pleiotropic and evolutionary constrained than younger Sfp genes

Gene properties that denote higher pleiotropy such as high level and breadth of expression are thought to be correlated with gene age ^25^. We investigated whether functional roles are evenly associated with the different age classes of Sfp genes. Only class A genes are enriched for GO terms (5%FDR) beyond those strictly associated with reproduction (*e*.*g*. sexual reproduction, insemination, or regulation of female receptivity), denoting a broader functional scope (Fig. 2a, Supplementary Table 3). This conclusion did not change when we repeated this analysis comparing Sfp genes originated before and after the radiation of the *Drosophila* genus (Supplementary Table 3). This augmented functional breadth is consistent with enhanced pleiotropy.

**Figure 2.**
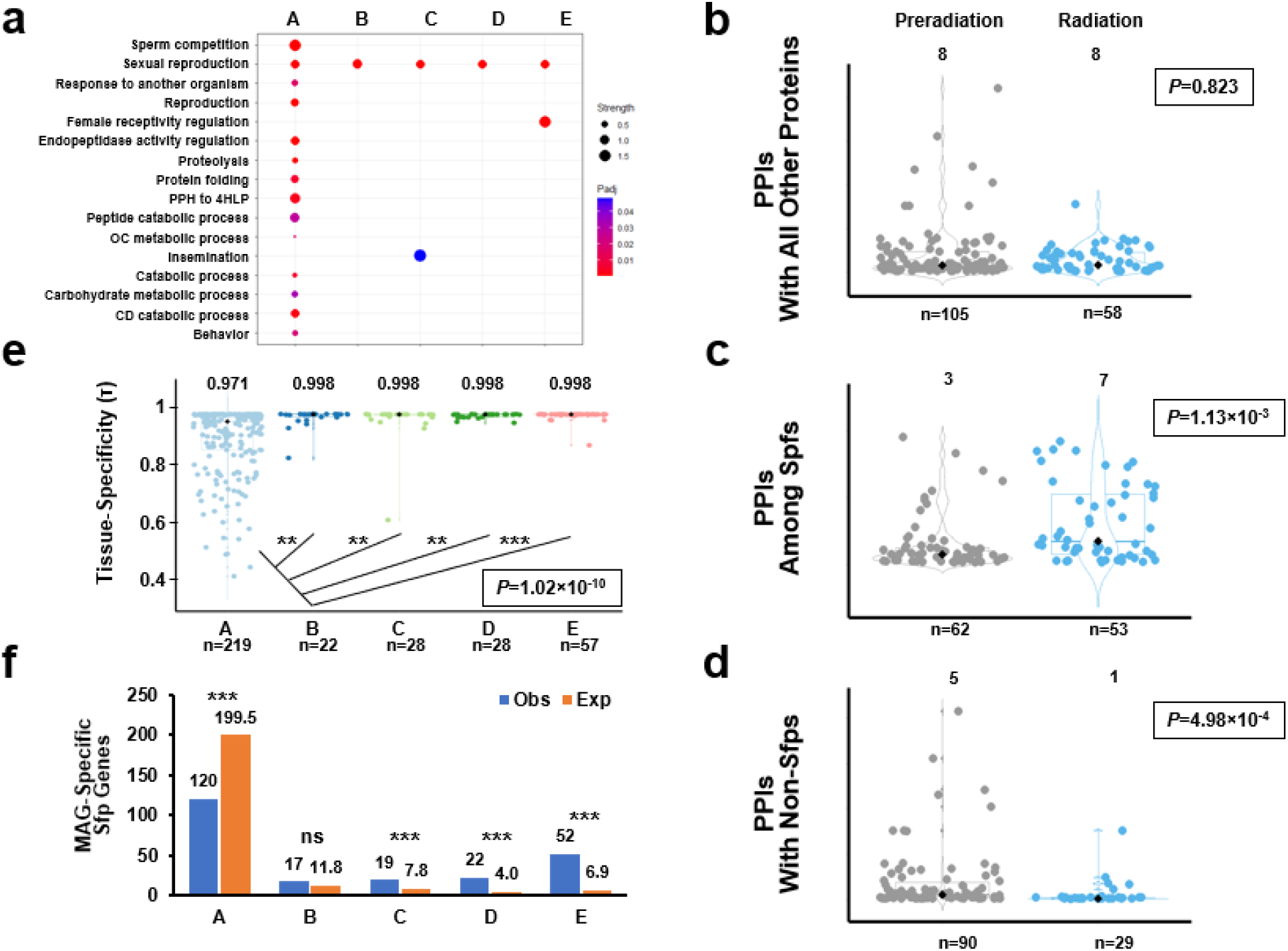
Ancient Sfps are more pleiotropic than younger Sfps. **a** Gene ontology enrichment based on FlyBase functional annotations followed by removal of redundant terms using REVIGO ^32^. The size of the dots in the dot plot are proportional to the gene-class enrichment and the dots are colored based on FDR-corrected p-values. **b-d** Violin and box plots showing the distribution of number of high-confidence protein-protein interactions (PPI) of the Sfps originated before and during the radiation of the *Drosophila* genus with other proteins, among Sfps, and with non-Sfps. **e** Violin and box plots showing the distribution of tau expression-specificity index of Sfp-encoding genes of different age classes based on FlyAtlas2 ^33^. **f** Clustered column plot comparing the observed (blue) and expected (orange) number of Sfp-encoding genes showing male reproductive gland (MAG) expression at τ≥0.9 across age classes. Expected counts were calculated based on the proportion of these age classes in the entire *D. melanogaster* gene complement ^17^. In violin plots, the *P* value associated with the Kruskal-Wallis rank sum test for differences across age classes is provided while the asterisks indicate the posthoc Wilcoxon rank-sum tests for which statistical significant difference were found after correcting for multiple tests ^34^. The median value for each distribution is shown on top and indicated with a black diamond. Whether the difference between observed and expected counts is statistically significant is indicated with asterisks according to the standardized residual values of the chi-square test of independence: *, <0.05; **, <0.01; ***, <0.001 ^24^.

Enhanced pleiotropy is thought to increase selective constraints, reducing evolutionary rates ^26,27^. However, studies examining pleiotropy’s effect on selection efficacy ^9,28-30^ have been inconsistent, partly due to conflictive results derived from interrogating different proxies of pleiotropy. We evaluated whether age class A genes showed enhanced pleiotropy by assessing different attributes. First, we retrieved reliable protein-protein interaction (PPI) data for 163 Sfp genes ^31^ (Supplementary Table 1). This includes interactions among Sfps and interactions with non-Sfps. One hundred and fifteen Sfp genes interacted with other Sfp genes, finding an equal representation of all age classes in the Sfp interactome (chi-square test of independence, χ^2^=6.35, d.f.=4, *P*=0.174). Examining protein connectivity across age classes showed no significant differences (Kruskal-Wallis rank sum test, χ^2^=5.06, d.f.=4, *P*=0.281) (Supplementary Fig. 2 and Supplementary Table 4), which holds when comparing the more broadly defined age classes, *i*.*e*. before and during the *Drosophila* radiation (*i*.*e*. A vs B+C+D+E; Wilcoxon rank-sum test, χ^2^=0.05, d.f.=1, *P*=0.823) (Fig. 2b). However, when we excluded interactions with non-Sfps and restricted our analysis to interactions among Sfp proteins, we found significant differences (Kruskal-Wallis rank sum test, χ^2^=14.255, d.f.=4, *P*=6.25×10^-3^), with age class A Sfps having significantly fewer interactions than the Sfps coded by genes in age class C, and a decreased number although not significant in relation to the Sfps from the age classes B and E (Supplementary Fig. 2 and Supplementary Table 4). The comparison between the pre- and during radiation age classes substantiates even further the difference in their degree of connectivity within the Sfp interactome (Wilcoxon rank-sum test, χ^2^=10.6, d.f.=1, *P*=1.13×10^-3^) (Fig. 2c). Notably, the analysis of interactions outside the Sfp interactome revealed age class A Sfps with significantly more interactions than other age classes (Kruskal-Wallis rank sum test, χ^2^=13.402, d.f.=4, *P*=9.5×10^-3^) (Supplementary Fig. 2 and Supplementary Table 4), which is again more obviously detected by the pre-vs during the radiation contrast (Wilcoxon rank-sum test, χ^2^=12.124, d.f.=1, *P*=4.98×10^-4^) (Fig. 2d). These results were consistent when considering only the 228 common Sfp genes (Supplementary Fig. 3).

**Figure 3.**
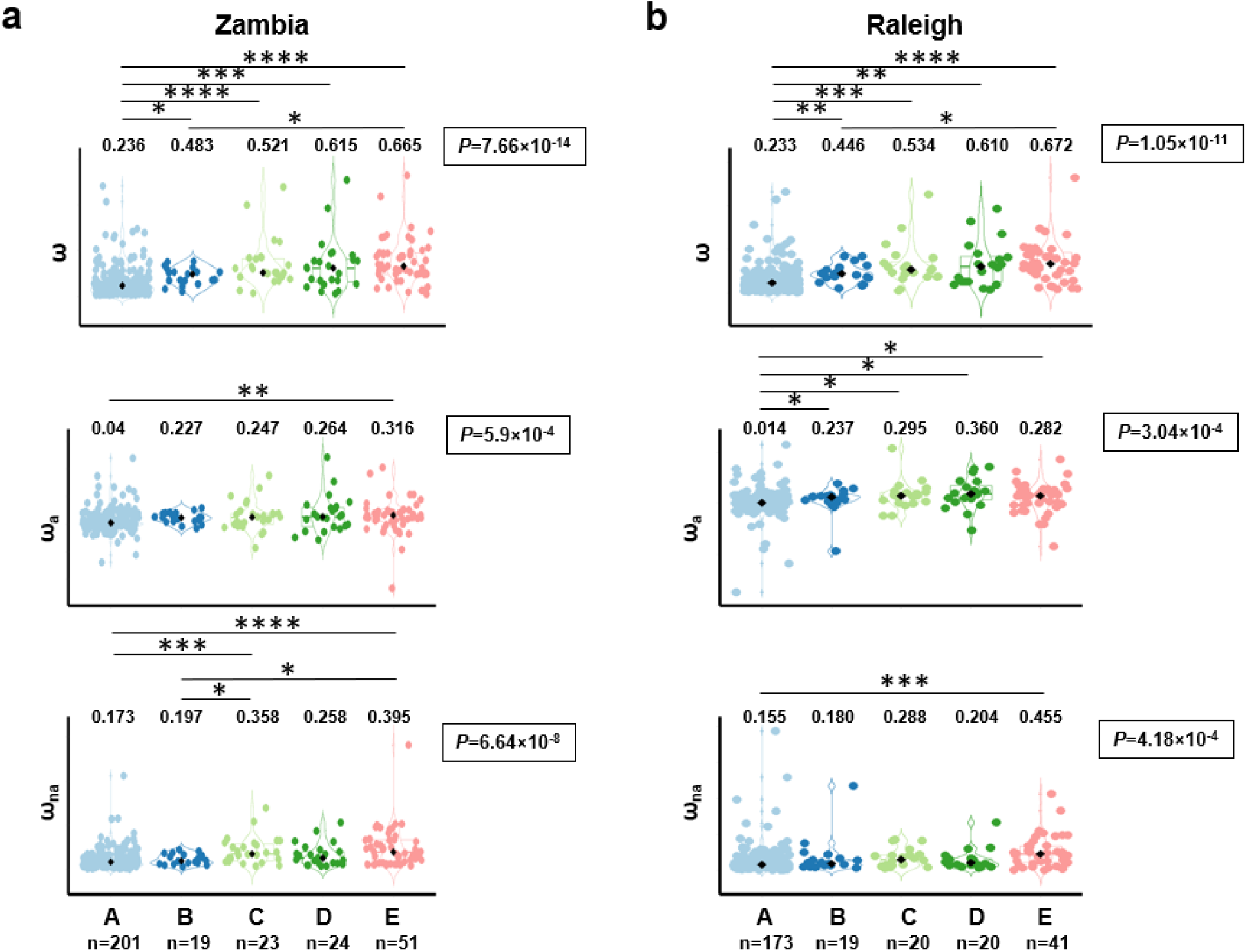
Distribution of coding sequence evolution metrics between *D. simulans* and two populations of *D. melanogaster* for Sfp genes across age classes. **a** The ratio of nonsynonymous to synonymous substitutions (ω), adaptive (ω_a_) and non-adaptive (ω_na_) evolution estimated using data from a *D. melanogaster* Zambia population. **b** Same estimates using data from a Raleigh population. Violin and box plots are provided for each age class by metric combination. The median value for each distribution is shown on top and indicated with a black diamond. The *P* value associated with the Kruskal-Wallis rank sum test for differences across age classes is provided per contrast. Lines on top, significant posthoc Wilcoxon rank-sum tests: *, <0.05; **, <0.01; ***, <0.001, ****, <0.001 ^24^.

Next, we investigated expression properties, hypothesizing that the broader functional range and interactive repertoire of class A Sfp genes would result in a broader expression profile. Indeed, class A genes have a significantly lower tau (τ) index compared to other age classes (Kruskal-Wallis rank sum test, χ^2^=52.62, d.f.=4, *P*=1.02×10^-10^) (Fig. 2e, Supplementary Tables 1 and 5), are less often male reproductive gland-specific (τ≥0.9 and maximum expression in that tissue; chi-square goodness-of-fit, χ^2^=425.51, d.f.=4, *P*<2.2×10^-16^) (Fig. 2f, Supplementary Table 1), and are in general less likely to show tissue-specificity (τ≥0.9 regardless of the tissue with maximum expression; chi-square goodness-of-fit, χ^2^=454.58, d.f.=4, *P*<2.2×10^-16^) (Supplementary Table 1), particularly in the case of the age class E. These results are recapitulated when only the 228 common Sfp genes are considered (Supplementary Table 5, Supplementary Fig. 4).

**Figure 4.**
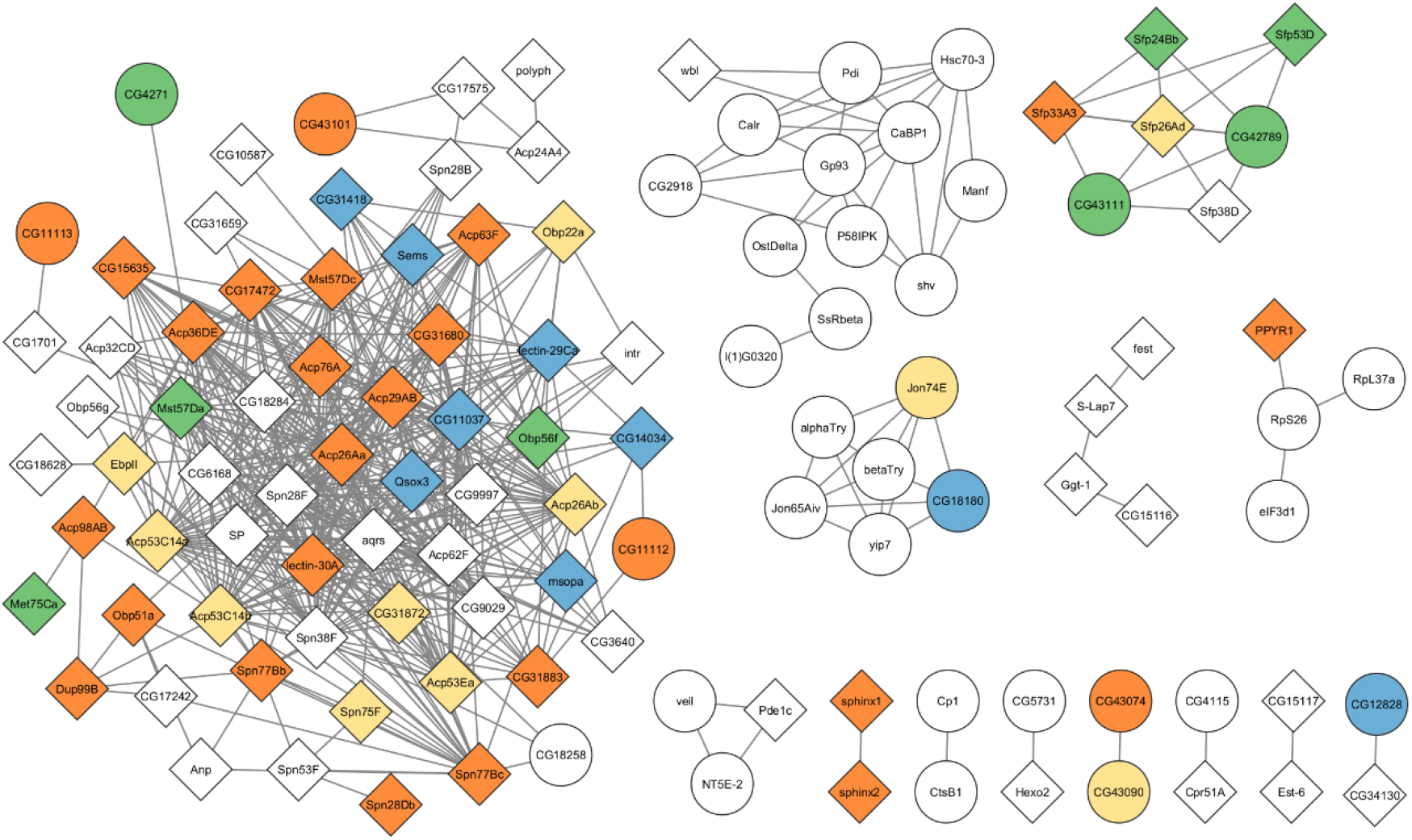
The Sfp protein-protein interaction network of *D. melanogaster*. The topology and composition of the Sfp interactome are shown. All 356 Sfp genes were considered. Only high-confidence interactions according to STRING (Methods) were included. Sfps with reproductive functions (*i*.*e*. those associated with the GO terms *sexual reproduction, reproduction, sperm storage, sperm competition, regulation of female receptivity, mating behavior, insemination*) are indicated with diamonds, while non-reproductive Sfps are shown by ovals. Age classes are color-coded: A (white), B (blue), C (yellow), D (green), and E (orange).

Lastly, we tested whether the increased pleiotropy of Sfp genes originated before the *Drosophila* radiation in terms of functional scope, protein connectivity, and expression breadth is associated with a slower rate of sequence coding evolution using inter- and intraspecific sequence information from the close relative *D. simulans* and two different *D. melanogaster* populations, one from Zambia (ZI) and one from Raleigh (RAL). Sfp genes in age class A show lower ratios (ω) of nonsynonymous to synonymous divergence, *i*.*e*. they evolve slower, than younger genes (RAL: chi-square goodness-of-fit, χ^2^=57.35, d.f.=4, *P*= 1.1×10^-11^; ZI: chisquare goodness-of-fit, χ^2^=67.50, d.f.=4, *P*=7.7×10^-14^) (Fig. 3). Differences in the rate of evolution can be influenced by both adaptive and nonadaptive evolutionary mechanisms. To appraise the effect of these two types of mechanisms on the rate of evolution of Sfps in relation to age, we used population polymorphism and divergence sequence data to estimate the proportion of substitutions fixed by positive selection (α) in order to derive estimates of adaptive (ωa) and non-adaptive (ωna) evolution, which provides a better understanding about how selection shapes gene evolution at the sequence level compared to using α alone ^35-37^. We found that the lower evolutionary rate (ω) of ancient Sfp gene is a consequence of constraints in both their adaptive (ωa) and non-adaptive (ωna) evolutionary rates compared to Sfps of younger age classes (Fig. 3, Supplementary Tables 6-7).

### The Sfp interactome includes subnetworks with distinct functional and evolutionary properties

The differences in the number of PPIs between age class A and younger age classes, regarding interactions with both Sfps and unrelated proteins, raise questions about their localization and aggregation within the Sfp interactome. Among the 163 genes with available PPI information, we found 98 forming six subnetworks comprising ≥4 members, with one dominant, core subnetwork containing 56% (64/98) of Sfps (Fig. 4, and Supplementary Table 1). The remaining 65 Sfps either did not form subnetworks above the minimum size or only interacted outside the Sfp interactome.

The core subnetwork expands as different age class Sfps are incorporated, being populated with many younger Sfps from the age class E (Supplementary Figs. 5-6). To quantify this and other trends, we assessed whether gene age is distributed evenly across subnetworks. The trend for age class E Sfp genes was not statistically significant (20/64, *P*_adj_=0.1687, Monte Carlo simulations, n=100,000) (Supplementary Table 8). This core subnetwork also includes all members, six, of the SP network with available PPI data ^11^. Further, the second- and third-largest subnetworks are significantly enriched for age class A and D Sfps, respectively (13/13, *P*_adj_=1.7×10^-4^, and 4/6, *P*_adj_= 6.8×10^-3^, respectively). Importantly, in line with our GO term enrichment analysis, the genes in the core subnetwork are significantly associated with reproductive roles, while the second and fourth largest subnetworks are depleted of this type of genes (chi-square test of independence, χ^2^=60.22, *P*_adj_=5.0×10^-4^, 2000 simulations; post-hoc tests, *P*_adj_<1.0×10^-6^ for the three age classes) (Supplementary Table 9).

Sfps are known to be among the fastest evolving at the sequence level between species ^38-41^. However, given the distinctive evolutionary dynamic of the core subnetwork, with younger genes enriched for reproduction functions and interactions with other Sfps, we predicted that Sfps in the core subnetwork would have different evolutionary rates relative to other Sfps. We found that the Sfp genes in core subnetwork evolved faster (Mann-Whitney; Raleigh: Z=-2.95, *P*=0.0031; Zambia: Z=-2.578, *P*=0.0099), with population-specific differences in adaptive (ωa) and non-adaptive (ωna) evolution rates (Supplementary Fig. 7 and Supplementary Table 10). In sum, these findings highlight unique functional and evolutionary characteristics of this relatively young reproductive subnetwork of the Sfp complement.

## CONCLUSION

Our findings illuminate the age-dependent nature of the functional and evolutionary characteristics of Sfp genes. Contrary to the prevailing belief that Sfp genes are rapidly evolving and young, we discovered a significant proportion of evolutionarily ancient Sfp genes. These genes have expanded their functional repertoire and connectivity while experiencing substantial evolutionary constraints. This discovery was made possible through refined genomic and functional data from several *Drosophila* species and related Diptera lineages. As similar data become available for other vertebrate and invertebrate species, we will be able to determine whether the functional and evolutionary trajectories of comparable Sfp gene complements are similar in lineages with different interactome architectures, population parameters such as the effective population size, or mating systems with distinct intensities of selection.

## METHODS

### Sfp gene complement

Two independent efforts have aimed to identify high-confidence Sfp-encoding candidate genes in *D. melanogaster* ^15,16^. With different emphasis in the type of evidence used, these studies consider transcriptomic and proteomic data, sequence homology searches, computational predictions of the signal peptide, and phenotypic data derived from gene perturbation studies. Due to the difficulty in reliably delineating a high-confidence Sfp gene set, it is likely that both attempts are affected by false positives and negatives until systematic molecular and phenotypic studies are implemented for each candidate gene. Therefore, we focused on 357 Sfps gene candidates deemed as high-confidence by either study, and repeated key analyses using a more conservative set of 228 genes deemed likewise by both publications (Supplementary Table 1).

### Phylogenetic gene age dating

Gene age inferences were based on syntenic alignments among 17 fly species plus *D. melanogaster* ^17^. These included two outgroup species, *S. lebanonensis* and *B. dorsalis*, from the Drosophilidae and Tephritidae families, respectively. A parsimony framework was used to assign genes to specific branches of the species phylogeny ^42^. Compared to previous attempts ^42,43^, the new information used by the authors incorporated genomes from more species, including 10 new, more contiguous genomes scaffolded with long PacBio HiFi reads, thus outperforming earlier ones utilizing Sanger sequencing reads ^44^. Excluding the *D. melanogaster* genome, all genome assemblies have a contig/scaffold N50 higher than 3 Mb.

### Sfp interactions and functional properties

RNAseq tissue expression values for 31 adult tissues and body parts from adult males and females were retrieved from FlyAtlas2 ^33^ and used to calculate the tissue specificity index (Tau Index, τ) (Yanai et al. 2005). This index takes values from 0 to 1, with higher values denoting a narrower expression profile. We deemed a gene as tissue-specific when its index value was at least 0.9. The number of protein-protein interactions (PPI) for each Sfp and the construction of the Sfp network was predicted with STRING v12 ^31^ under the high confidence threshold and excluding text mining scores. The PPI network was visualized with Cytoscape v3.10 ^45^. Enrichment for biological process Gene Ontology terms was examined within STRING at a 5% FDR ^34^ and redundant terms were excluded using REVIGO ^31^.

### Rates of nucleotide sequence evolution

Synonymous and nonsynonymous nucleotide substitutions for each gene were retrieved from sequence comparisons between *D. simulans* and 197 lines from an African (Zambia, ZI) as well as 205 lines from a North American (Raleigh, RAL) population of *D. melanogaster* ^46-48^ using the iMKT R package ^49^. This package provides single-gene estimates for different population genetic parameters, relying in alignment pipelines and filtering criteria described in detail elsewhere ^47,50^, including divergence statistics derived from curated alignments between the two species ^51^. From iMKT, we downloaded the allele frequency spectrum and applied a 5% frequency threshold to estimate the number of neutral segregating sites in the non-synonymous class ^49^. These corrected estimates were used to calculate α, the proportion of nucleotide substitutions driven by positive selection ^49,52^, which was used in turn to calculate the rates of adaptive (ωa) and non-adaptive molecular evolution (ωna) ^36,37^.

### Statistical analyses

Two-tailed Fisher’s exact test, chi-square goodness-of-fit, chi-square test of independence, analysis of residuals, Kruskal-Wallis rank sum, Mann-Whitney pairwise tests, and the Benjamini-Hochberg correction for multiple testing were conducted using built-in functions in R ^53^. Monte Carlo simulations (n=10,0000) were performed by resampling without replacement.

### Declaration of Generative AI and AI-assisted technologies in the writing process

During the preparation of this manuscript the authors used ZotGPT (GPT-4 Omni) to improve readability and language. After using this tool, the authors reviewed and edited the content as needed and take full responsibility for the content of the publication.

## DATA AVAILABILITY

All data are available in the supplementary files.

## CODE AVAILABILITY

Code for Monte Carlo simulations is available on GitHub (https://github.com/Ranz-Lab/SFPsimulations) and Zenodo ^54^.

## ACKNOWLEDGEMENTS

We thank Manyuan Long for granting us early access to unpublished gene age data, and Sara Good, Wilfried Haerty, and Rama Singh for comments on the manuscript. This work was supported by a Natural Science Foundation grant to J.M.R (MCB-2129845), and a NSERC Discovery Grant (RGPIN-2017-04599) to A.C.

## AUTHORS CONTRIBUTIONS

**Alberto Civetta**, Data curation, Formal analysis, Investigation, Methodology, Writing – original draft, Writing – review and editing; **Carolina Flacchi**, Data curation, Formal analysis; **Imtiyaz E. Hariyani**, Formal analysis; **Jose M. Ranz**, Data curation, Formal analysis, Investigation, Methodology, Writing – original draft, Writing – review and editing.

## COMPETING NTERESTS

The authors declare no competing interests.

